## Supplementary Figures for "RNA tertiary structure and conformational dynamics revealed by BASH MaP"

**­**
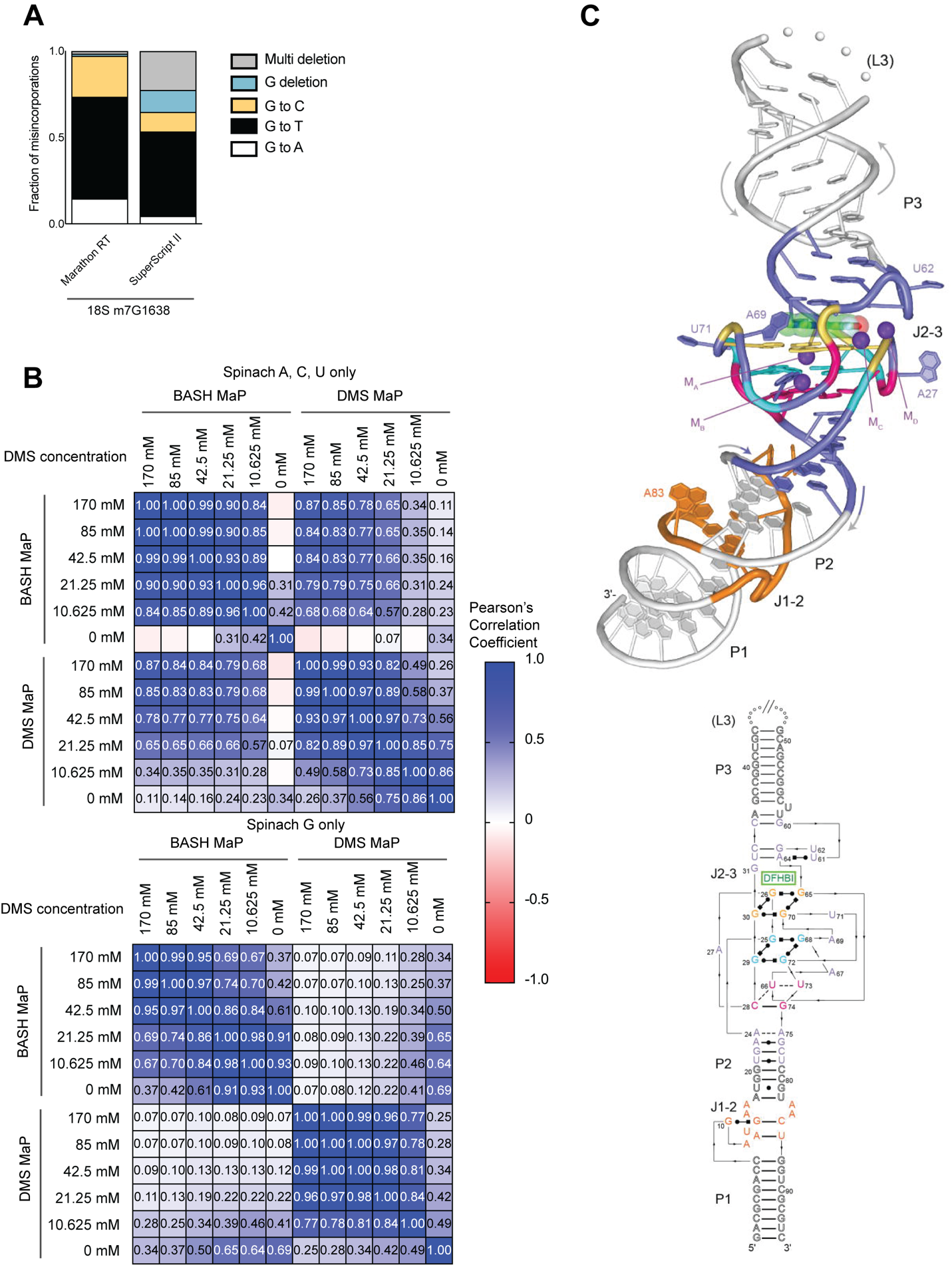

**Supplementary Data Fig. 1 | Comparison of reverse transcriptases on the misincorporation signature of abasic sites and BASH MaP reproducibility and correlation with DMS MaP**

(**A**) The misincorporation signature of abasic sites generated by Marathon RT exhibits fewer deletions than SuperScript II. In **Fig. 1D**, we examined the misincorporation signature of G’s after DMS MaP and BASH MaP with SuperScript II and found high rates of deletion. We therefore asked if the high deletion rate is due to the choice of reverse transcriptase or if it is an intrinsic property of abasic sites during reverse transcription. The mechanism of deletion is important because deletions must be ignored in further data processing steps as these misincorporation are often unable to align to a specific guanine if there are two or more consecutive guanines. We chose to test Marathon RT as this reverse transcriptase was previously shown to induce lower deletion rates than SuperScript II^1^. As shown, Marathon RT produces predominantly G🡪T transition and with an overall much lower rate of deletions during reverse transcription of abasic sites. Thus, the deletions seen in SuperScript II BASH MaP data are likely the result of features unique to the SuperScript II enzyme. Future implementations of BASH MaP may benefit from using Marathon RT.

(**B**) BASH MaP and DMS MaP display high reproducibility and show high correlation for A, C, and U bases. In **Fig. 1F-G**, we establish the reproducibility of BASH MaP and show that misincorporation data at A, C, and U bases is not compromised by BASH treatment. Here, we exhaustively examine the reproducibility and correlation between BASH MaP and DMS MaP protocols. Spinach RNA was first probed with the indicated concentrations of DMS for 8 min at 25°C using bicine (200 mM) pH 7.75 buffer conditions^2^. Then, DMS-treated RNA was split into two fractions. One fraction underwent BASH treatment followed by reverse transcription (BASH MaP), while the other fraction was immediately reverse transcribed with SuperScript II (DMS MaP). We then sequenced the RNA, calculated the misincorporation rate for each nucleotide, and calculated a Pearson’s correlation coefficient between each sample. A, C, and U nucleotide data was analyzed separately from G data. The heatmap of correlation coefficients shows that both BASH MaP and DMS MaP display high reproducibility over a range of DMS concentrations. Furthermore, BASH MaP and DMS MaP display strong correlations for A, C, and U misincorporation data at higher DMS concentrations. As expected, DAGGER MaP and DMS MaP display no correlation among G misincorporation data.

(**C**) Spinach crystal structure 4TS2^3^. Top, Spinach crystallographic structure. Bottom, partial secondary structure schematic of the Spinach crystal structure with annotated domains. G-quadruplex G’s are indicated below in orange and blue. The mixed tetrad is indicated below in pink.

**­**
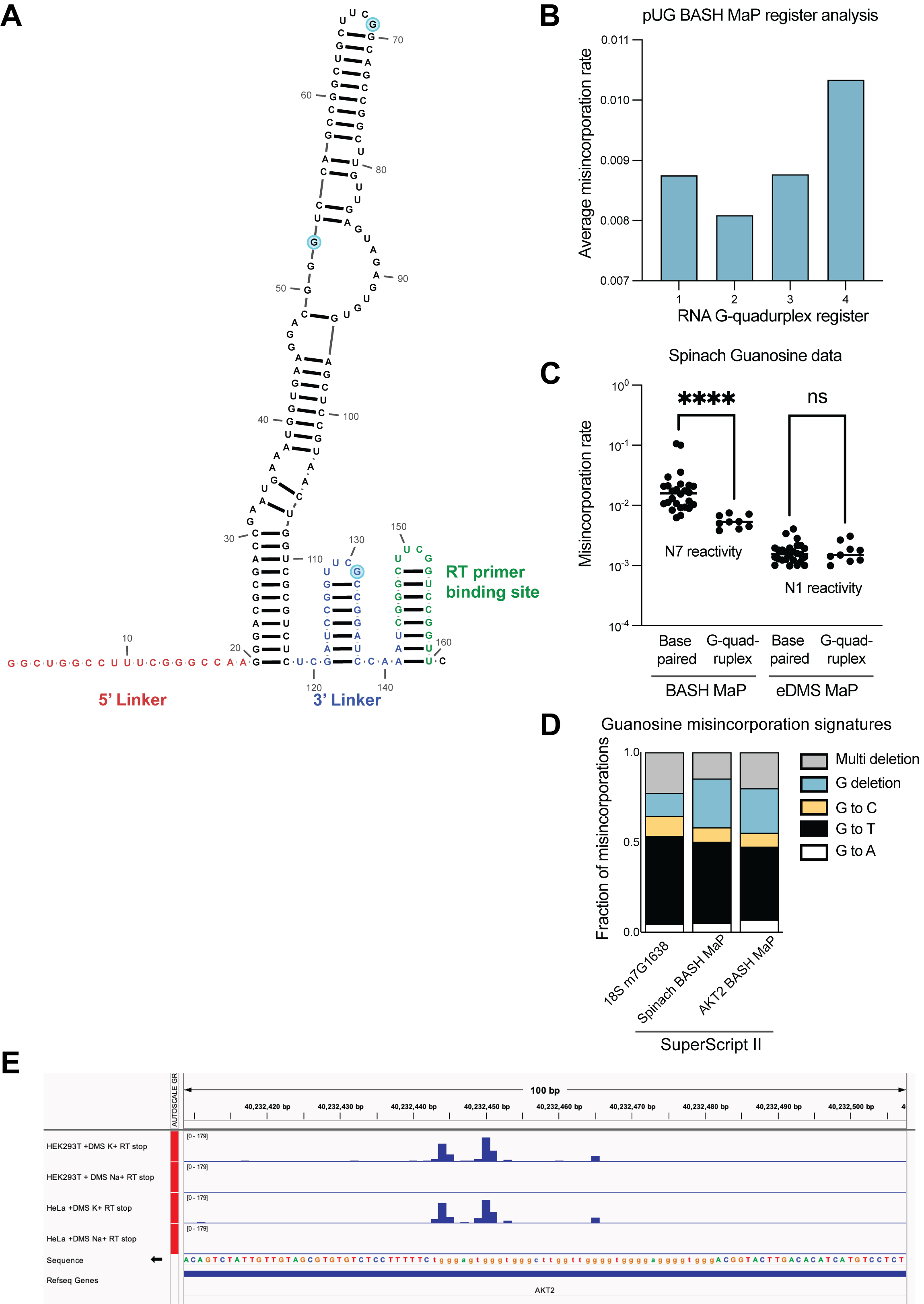
**Supplementary Data Fig. 2 | DMS MaP is unable to efficiently detect accessibility of the N7 position in guanine and validation of *AKT2* N7G protections in a detected G-quadruplex.**

(**A**) Secondary structure of Spinach RNA model used for in vitro probing experiments. To aid with DMS/BASH MaP library preparation, Spinach was in vitro transcribed with an additional 5’ linker sequence (red) as well as with an additional 3’ linker (blue) and RT primer binding site (green). Due to the use of PCR in library preparation, misincorporation data is unavailable for the 5’ linker and RT primer binding site in the utilized Spinach construct. Three single-stranded G’s in the Spinach structure cassette are highlighted in light blue. The secondary structure model for Spinach was derived from the crystal structure (PDB: 4TS2) and linker regions were folded using mFOLD^4^.

(**B**) Conformational analysis of alternative 15x polyUG G-quadruplex registers. In **Fig 2.E-F**, we found that different G-quadruplex registers can be differentiated by N7G reactivity data. It seemed likely that the 15x polyUG repeat RNA populates each register with equal stoichiometry. To test this, we reasoned that if each register were equally populated, then the average misincorporation rate of the G’s engaged in a G-quadruplex for each register should be the same. Therefore, we asked whether the average misincorporation rate of the G-quadruplex G’s for each register was the same. Surprisingly, we found that Register 2 displayed the lowest average misincorporation rate among the four registers which suggests that Register 2 is populated with a higher stoichiometry than the other three registers.

(**C**) Comparison of BASH MaP and DMS MaP for discrimination of base-paired G’s versus G’s engaged in a G-quadruplex. Chemical probes that react with the Watson-Crick face of G have previously been used to assess the folding state of G-quadruplexes^5^. We reasoned that this strategy to profile G-quadruplex folding is prone to false positives because chemical probes that react with the Watson-Crick face, i.e., the N1 position of G, should show reduced chemical reactivity both when a G is base paired and when a G is in a G-quadruplex. Therefore, we asked whether DMS MaP optimized to methylate and detect N1G adducts (eDMS MaP) could differentiate between base-paired G’s and G-quadruplex G’s^6^. To test this, we probed Spinach with DMS (170 mM) for 6 min at 37°C using bicine (200 mM) pH 8.37 buffer conditions to promote the formation of m^1^G. To specifically detect m^1^G adducts, we utilized the property of SuperScript II to specifically encode m^1^G adducts as G🡪T and G🡪C misincorporations. We then prepared a BASH MaP library from the same DMS-treated Spinach sample and compared the two methods for their ability to differentiate G’s engaged in a G-quadruplex. We assigned G’s to either base-paired or G-quadruplex groups, plotted misincorporation rates and performed a Mann-Whitney U test to determine if the groups were statistically different from each other. On the left, BASH MaP, which measures N7 reactivity, clearly differentiates G-quadruplex G’s from base-paired G’s. On the right, eDMS MaP, which measures N1 reactivity, was unable to differentiate G-quadruples G’s from base-paired G’s. Taken together, this data shows that BASH MaP uniquely discriminates G’s engaged in G-quadruplexes from G’s engaged in base-paired interactions. ********p < 0.0001, ns p=0.9852, Mann-Whitney U test.

(**D**) Misincorporation signature of G’s in BASH MaP of *AKT2*. We wanted to validate whether misincorporation data at G’s in BASH MaP of *AKT2* produced misincorporations consistent with methylation at N7G. To test this, we compared the misincorporation signature of all G’s in *AKT2* with the misincorporation signature of G’s in BASH-treated Spinach and at the 18S m^7^G1638 site. All three experiments utilized SuperScript II as the reverse transcriptase. Comparison of misincorporation signatures for the three experiments revealed a common G🡪T and G deletion misincorporation signature. Consistent misincorporation signatures indicates that misincorporations at G’s in *AKT2* represent N7G methylation events.

(**E**) *AKT2* 3’UTR contains an in cellulo folded G-quadruplex. We wanted to test whether BASH MaP could identify N7G sites with low reactivity in cells. To identify a candidate G-quadruplex that is likely to be folded in cells, we turned to a previously published study which measured in cell G-quadruplex folding^7^. This study used a method that combined in cell-DMS treatment and subsequent RT stops. In brief, the in-cell DMS treatment methylates all N7 positions not engaged in a G-quadruplex. Then, cellular RNA is purified before undergoing refolding and RT stop profiling in K+ vs Na+ buffer. The presence of RT stops in K+ buffer implies that a G-quadruplex was present in the cell, protected from DMS, and able to refold into a G-quadruplex in the RT step. Shown are potassium-dependent RT stops in both HeLa and HEK293T cells. Thus, this region of the *AKT2* 3’UTR contains a folded G-quadruplex in cells, of an unknown topology or conformation. The presence of multiple RT stop sites, which occur at the 3’ end of G-quadruplexes, suggests that this region adopts multiple G-quadruplex conformations with different 3’ ends.

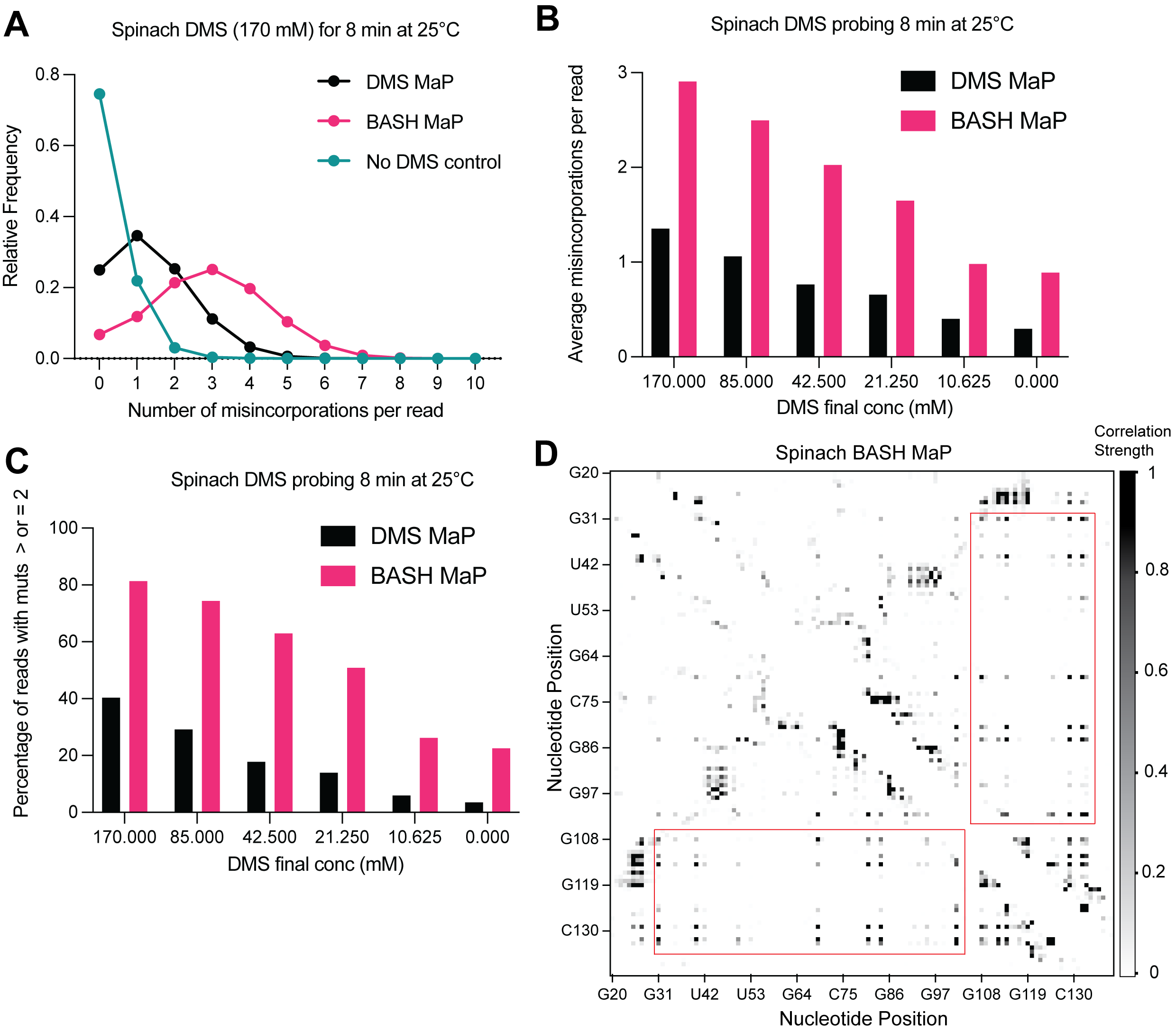
**Supplementary Data Fig. 3 | BASH MaP improves single molecule analysis through increased misincorporation density**

(**A**) Histogram of misincorporation per sequencing read reveals increased misincorporation density in BASH MaP. Single molecule analysis of structure probing data relies on the presence of multiple misincorporations per sequencing read. The number of misincorporations per sequencing read has been limited by the ability to simultaneously detect multiple methylations on a single RNA molecule. Since BASH MaP enables detection of N7G methylation events, we sought to quantify the increase in misincorporations per sequencing read after BASH treatment. We analyzed data from paired DMS MaP and BASH MaP datasets produced from the same pool of DMS-modified RNA and plotted a histogram representing the number of misincorporation per sequencing read. For reference, a full-length sequencing read for the Spinach RNA was 141 nucleotides long. The histogram revealed that the most common sequencing read for the no DMS-modified RNA had zero misincorporation. In contrast, the most common sequencing read for DMS MaP contained one misincorporation whereas the most common sequencing read for BASH MaP contained three misincorporations. Together, this shows that the most common sequencing read for BASH MaP enables observation of three times the number of DMS-methylation events as compared with DMS MaP.

(**B**) Plot of misincorporations per read shows BASH MaP doubles the misincorporations per read as compared to DMS MaP. To further identify whether BASH MaP data is suitable for single molecule analysis, we analyzed the average number of misincorporation per read for a range of DMS concentrations. The plot shows that the number of misincorporation per read is dependent on both DMS concentration and BASH treatment. BASH MaP samples roughly had twice as many misincorporation per sequencing read as DMS MaP libraries prepared from the same RNA. These results suggests that BASH MaP is suitable for single molecule analysis.

(**C**) Quantification of **Supplementary Data Fig. 3B** shows BASH MaP doubles the number of sequencing reads which can be used for single molecule analysis. Since single molecule analysis requires two or more misincorporation per read, we quantified the percentage of sequencing reads which meet this criterion for various DMS concentrations. A plot of the percentage of reads which could be used for single molecule analysis reveals that BASH MaP on average doubles the percentage of sequencing reads with two or more misincorporation when compared with paired DMS MaP datasets.

(**D**) Base-paired G’s display low levels of correlated mutations with other base-paired guanines. A heatmap of normalized correlation strength for all possible base-pair combinations in BASH-treated Spinach reveals a checkerboard-like pattern (red boxes). These points correspond to base-paired G’s which appear to co-mutate with other base-paired G’s.

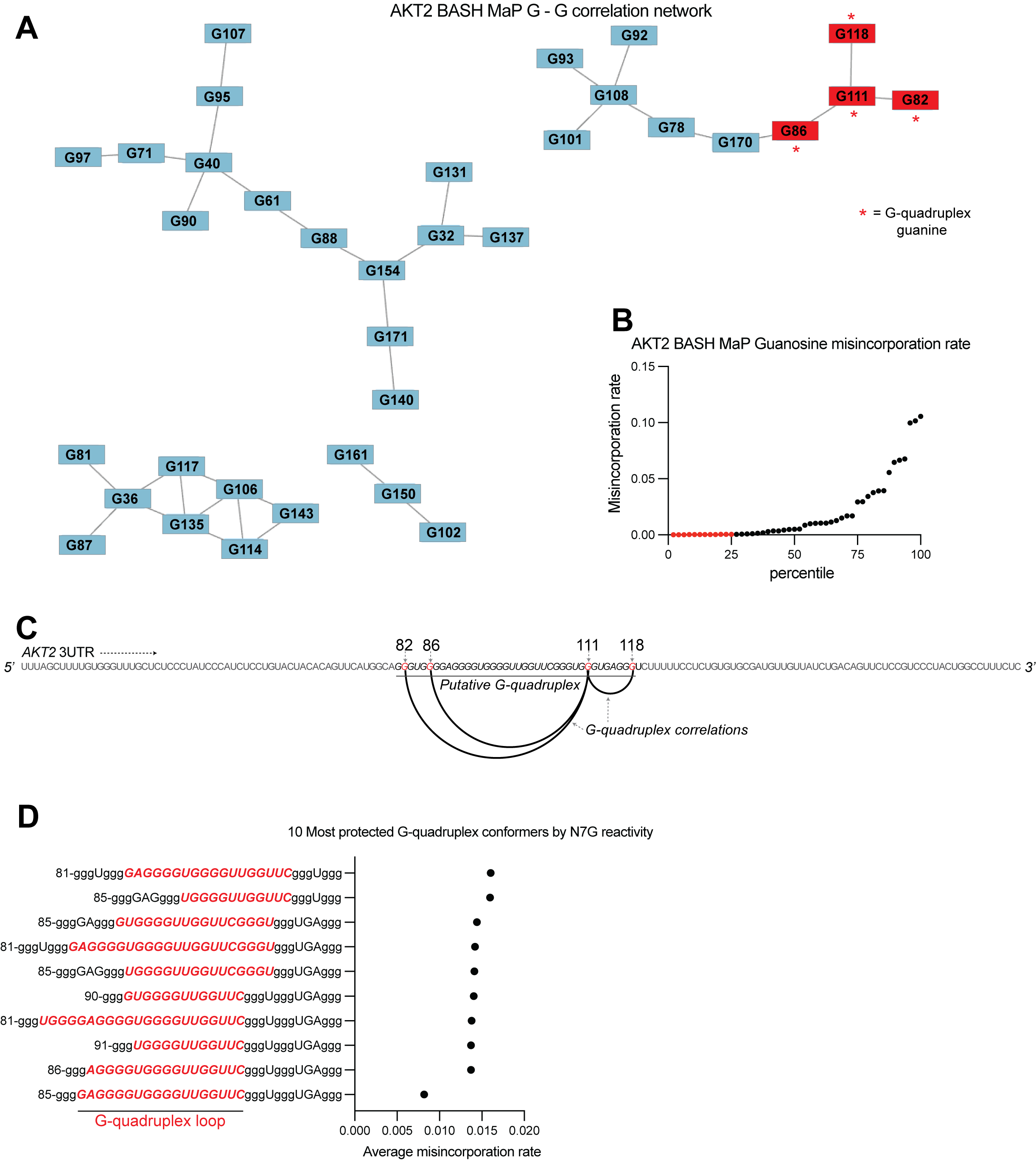
**Supplementary Data Fig. 4 | *AKT2* 3’UTR adopts an atypical G-quadruplex with a long central loop.**

(**A**) G – G correlation network for BASH MaP of *AKT2* 3UTR. In **Fig. 3G**, we show that network analysis of G – G correlations in Spinach BASH MaP data visualizes the G-quadruplex. We next wanted to know if a similar network analysis could identify a G-quadruplex in the *AKT2* 3’UTR. Like Spinach prior to its crystallographic structure determination, it is unclear which G’s are engaged in a G-quadruplex, especially since *AKT2* displays numerous G’s with low N7G reactivity. The correlation network for *AKT2* BASH MaP displays four clusters of G’s. Lowly reactive G’s as identified in **Supplementary Data Fig. 4B** display a cluster of four G’s, marked with red asterisks, which appear to be engaged in a G-quadruplex.

(**B**) *AKT2* 3’UTR N7G reactivity filtering. In **Fig. 5C**, we describe the criteria for identifying a G as engaged in a tertiary interaction. The first step in this process involves identification of the bottom quartile of G’s for N7G reactivity shown as red dots. We then applied this filtering process to the population average reactivity data for *AKT2* BASH MaP. From these G’s, we then identified G-quadruplex G’s which were both in the bottom quartile and connected in the network plot and marked these connections with red asterisks in **Supplementary Data Fig. 4A**.

(**C**) Arc plot of G-quadruplex G – G correlations in *AKT2* BASH MaP data. We first asked whether the G-quadruplex-correlated G’s, as identified in **Supplementary Data Fig. 4A**, could give insight into which G tracts were likely to be engaged in a G-quadruplex. Interestingly, an arc plot depicting the correlations between identified G-quadruplex G’s suggests that a major conformation of the *AKT2* 3’UTR G-quadruplex consists of the first two and last two guanine tracts. This finding is surprising because this G-quadruplex conformation would involve a large central loop which is uncommon for G-quadruplexes^8^.

(**D**) Conformational analysis of *AKT2* 3’UTR G-quadruplex suggests a large central loop. Since the *AKT2* 3’UTR sequence contains seven tracts of three or more guanines, we reasoned that this region may adopt many unique G-quadruplex conformations. This is supported by RT stop profiling which identified at least three distinct 3’ ends of various G-quadruplex conformations (**Supplementary Data Fig. 2F**). To calculate all possible three-tiered G-quadruplex conformations we utilized the program QGRS Mapper^9^. This software identified 129 unique three-tiered G-quadruplex conformations capable of forming in the *AKT2* 3’UTR. We then asked which conformations were most consistent with the population average N7G reactivity data. We reasoned that we could identify the most populated conformations by calculating the average misincorporation rate of the G-quadruplex G’s for each unique conformation. The conformations with the lowest average misincorporation rates are therefore likely to be the most populated conformations. We calculated the average misincorporation rate of the G-quadruplex G’s and plotted the ten conformations with the lowest reactivity. The notation in the plot is as follows: the first number indicates the number of the first nucleotide in the G-quadruplex sequence. G’s engaged in a G-quadruplex are identified as a lowercase ‘g’. The ten G-quadruplex conformations with the lowest N7G reactivity revealed a common large central loop, highlighted in red. Together, these data further support a model in which the *AKT2* 3’UTR adopts an unusual G-quadruplex conformation with a long central loop.

**­­**
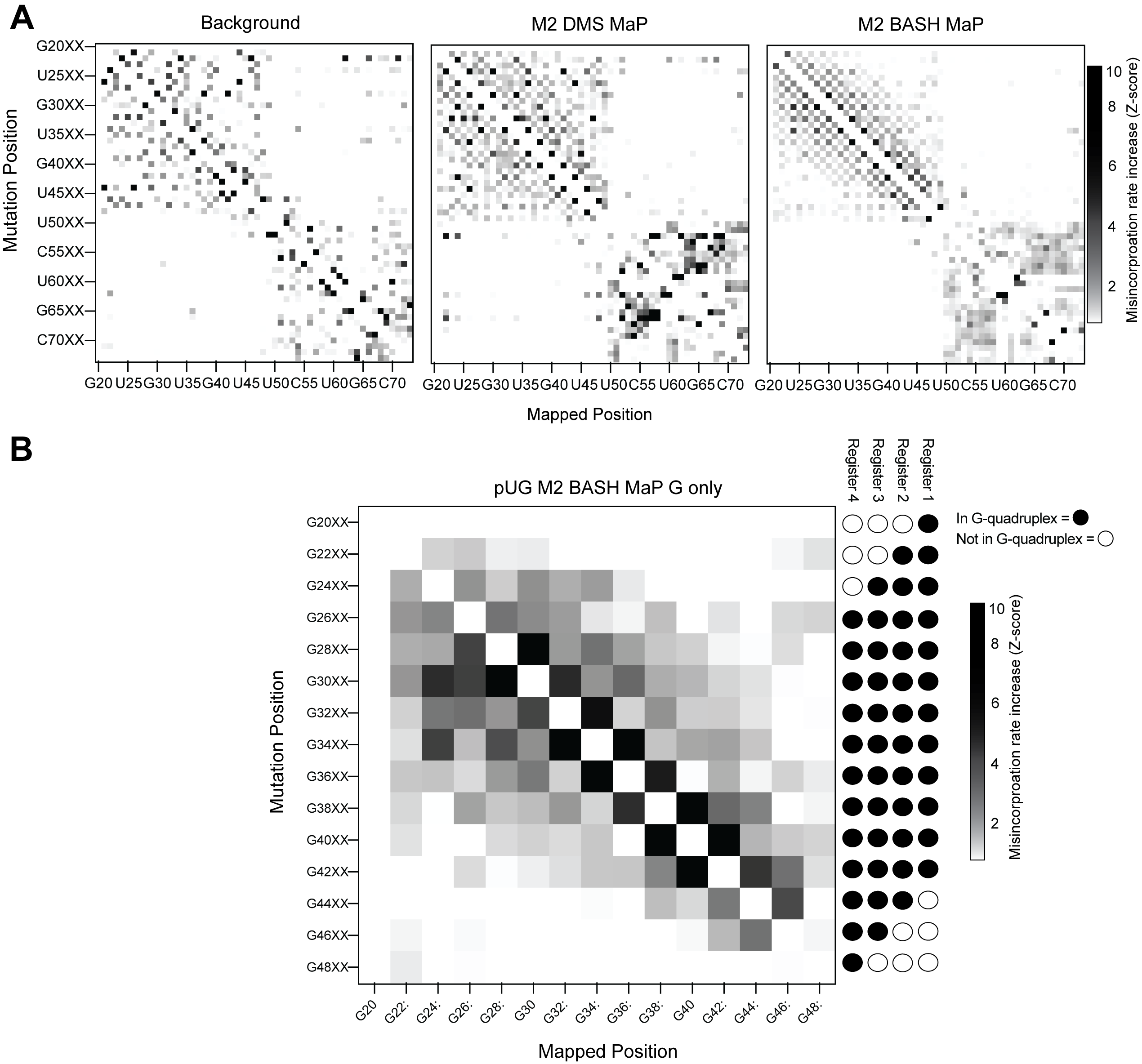
**Supplementary Data Fig. 5 | M2 BASH MaP of 15x polyUG repeat RNA**

(**A**) M2 BASH MaP of 15x polyUG repeat RNA reveals mutation-induced G-quadruplex unfolding. We next asked whether PCR-derived point mutations destabilized the G-quadruplex in the 15x polyUG repeat RNA. To test this, we performed random mutagenesis with PCR (left) and then performed M2 DMS MaP (middle) or M2 BASH MaP (right). Like the heatmap of M2 BASH MaP of Spinach in **Fig. 4B**, M2 BASH MaP of the polyUG RNA displayed a strong checkerboard pattern which occurred at pairs of G’s. As seen in Spinach, mutation of a G in a G-quadruplex appears to induce increased N7G reactivity of adjacent G’s.

(**B**) M2 BASH MaP of 15x polyUG repeat RNA supports a four-register model of RNA conformation. To validate whether mutations in the G-quadruplex cause global destabilization of the 15x polyUG repeat RNA, we analyzed how mutations at G-quadruplex G’s affected N7G reactivity of all other G’s. As expected, mutations in G’s that only populate one or two G-quadruplex registers had minimal impact on the global N7G reactivity. In contrast, mutations at G’s expected to populate all four G-quadruplex registers induced global increases in N7G reactivity. Mutations at G’s expected to populate only three G-quadruplex registers only induced local increases in N7G reactivity. The M2 BASH MaP heatmap for the 15x polyUG repeat RNA further supports the existence of alternative G-quadruplex registers and suggests that point mutations cause global destabilization and unfolding.

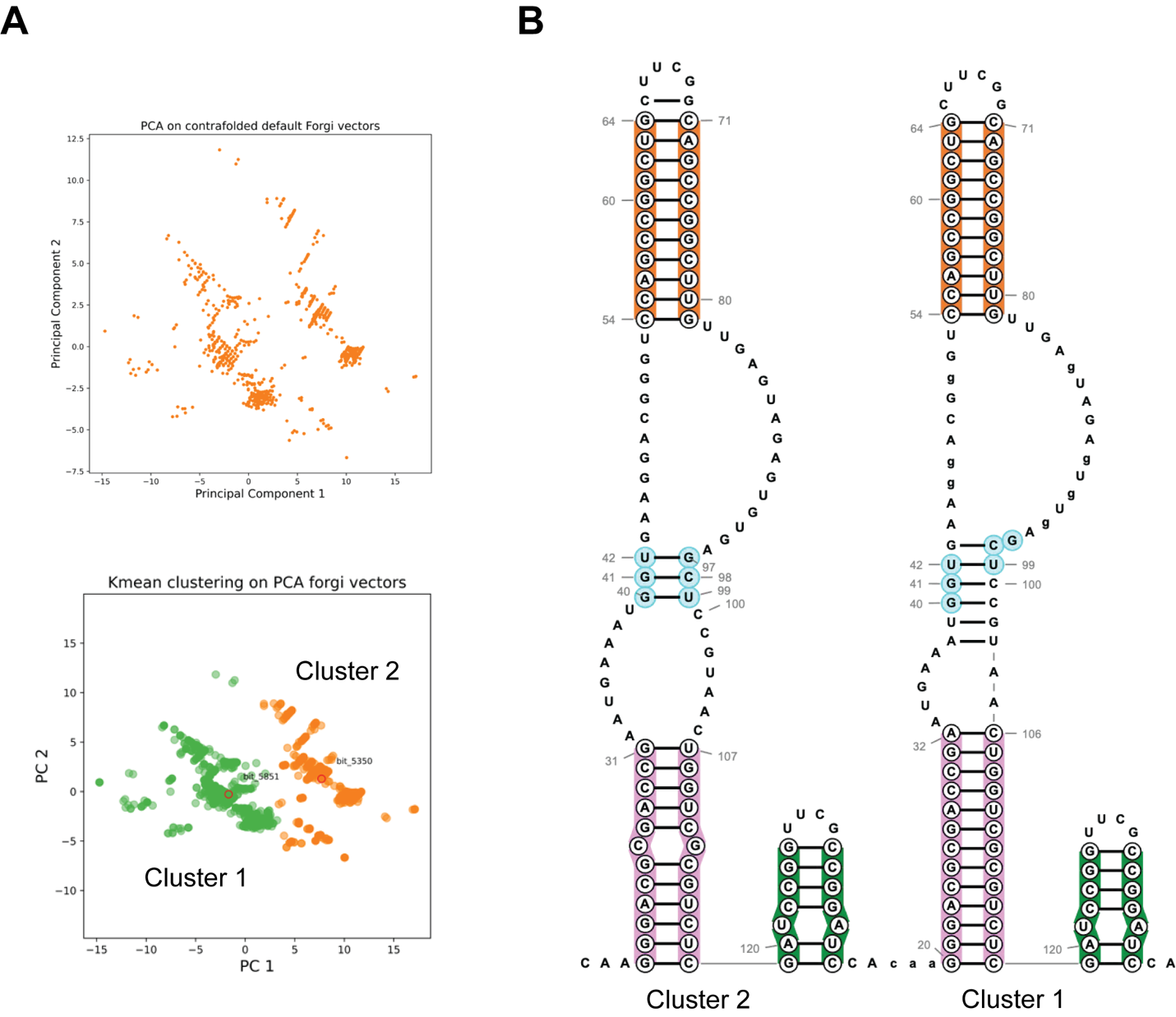
**Supplementary Data Fig. 6 | DAGGER clustering of Spinach BASH MaP**

(**A**) DAGGER clustering of 10,000 reads for BASH-MaP-treated Spinach. Principle component analysis revealed roughly two major clusters of RNA conformation. Kmeans clustering was used to identify the most representative secondary structure for each of the clusters. Cluster 1 is colored green. Cluster 2 is colored orange.

(**B**) Comparison of Cluster 1 and Cluster 2 representative secondary structures reveals alternative base pairing in P2 domain. Enlarged image of **Fig. 6I** depicts overall conservation of P1 and P3 domains between the two clusters. Differences between Cluster 1 and Cluster 2 are depicted in the alternative base pairing pattern in the P2 stem, colored light blue.

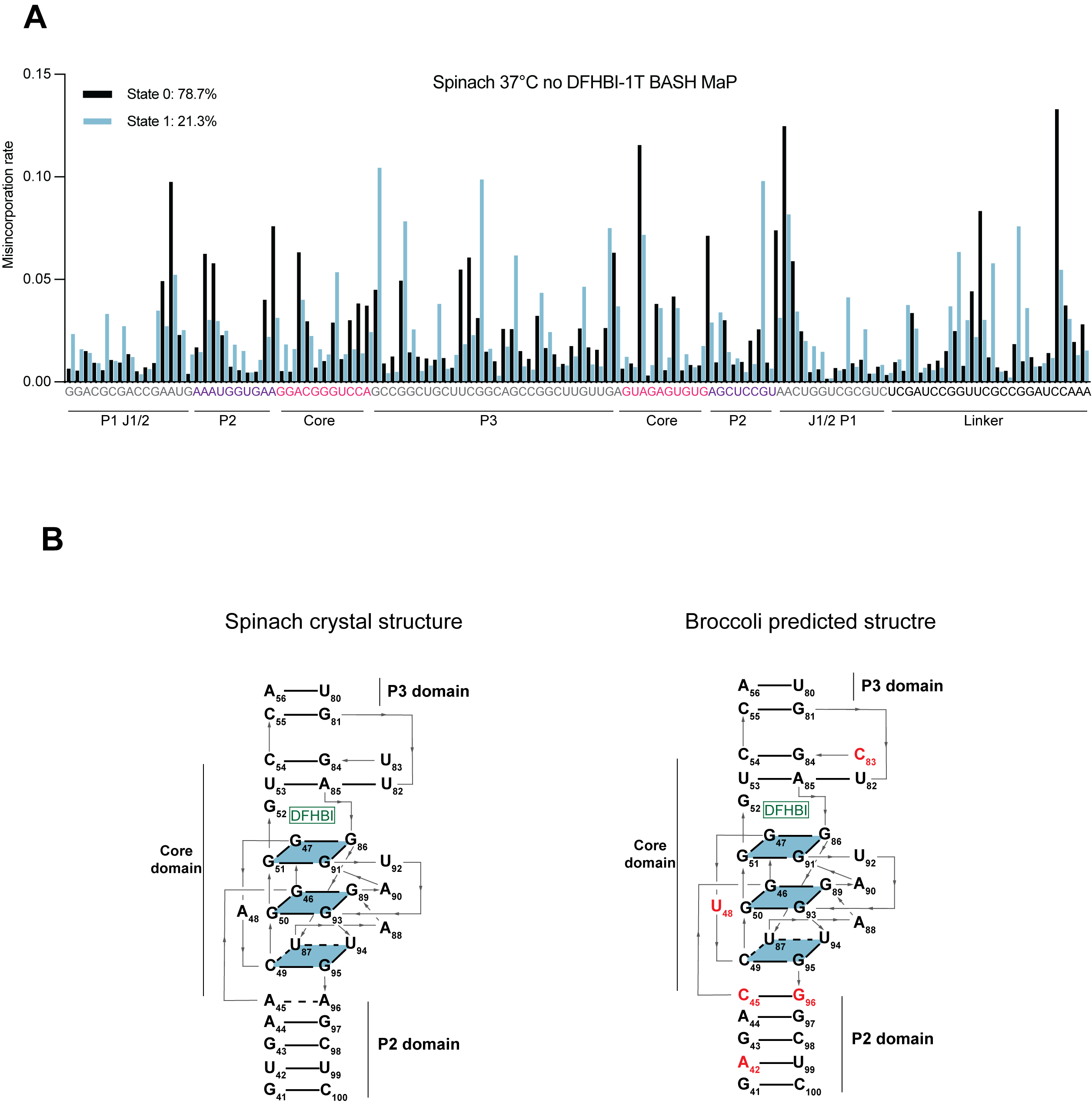
**Supplementary Data Fig. 7 | DANCE clustering of Spinach BASH MaP data and comparisons to Broccoli**

(**A**) DANCE deconvolution of BASH MaP treated Spinach identifies two conformations. To validate that the DANCE deconvolution of M2 BASH MaP was not due to PCR-derived mutations, we performed BASH MaP on non-mutagenized Spinach at an elevated temperature and without its ligand DFHBI-1T. DANCE clustering identified two conformations with similar abundances to **Fig. 6A**. As seen in the DANCE conformations of M2 BASH MaP, State 0 is consistent with a folded G-quadruplex whereas State 1 suggests a misfolding of the G-quadruplex in the core domain. These results suggests that the DANCE-identified clusters represent true alternative conformations of Spinach and are reproducible under a range of probing conditions.

(**B**) Alignment of Spinach and Broccoli three-dimensional structures. The predicted Broccoli three-dimensional structure displays an identical G-quadruplex topology to Spinach. Nucleotide differences in the Broccoli structure are indicated in red. Broccoli structure numbering was adjusted to align with the Spinach numbering. Key differences involve the strengthening of a C-G-C base triple above the ligand binding pocket, a change of a bulge A between the tetrad and a G-quartet to a U and three nucleotide differences in the P2 domain.

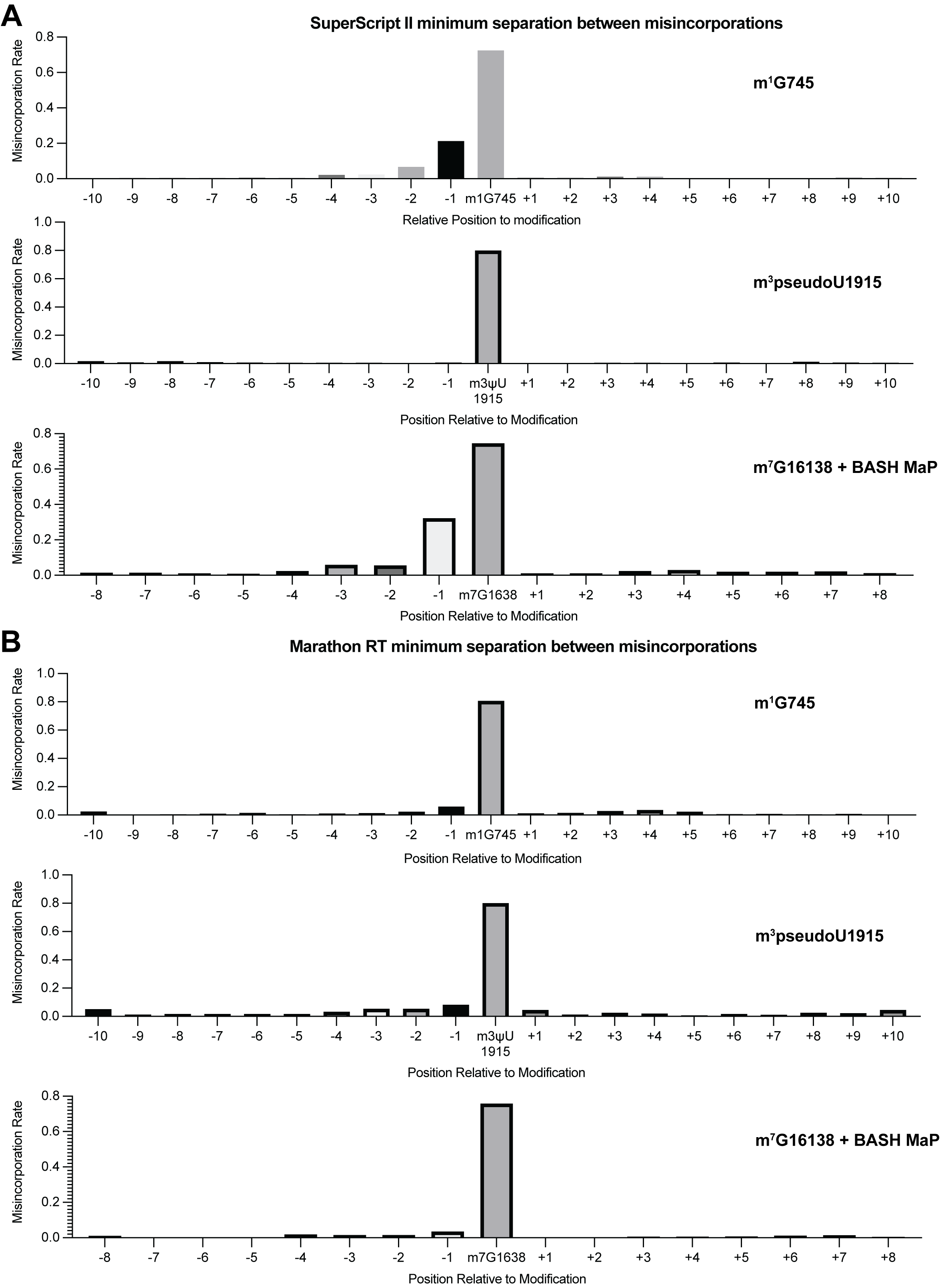
**Supplementary Data Fig. 8 | Validation of minimum separation between misincorporations for SuperScript II and Marathon RT**

(**A-B**) Misincorporations induced by SuperScript II (**Supplementary Data Fig. 7A**) and Marathon RT (**Supplementary Data Fig. 7B**) at m^1^G, m^3^psuedoU, and abasic sites. Misincorporations in DNA sequencing can be simple such as a G🡪T transition. Previous studies found that certain reverse transcriptases produce more complex types of misincorporations such as clusters of misincorporations for a single modified nucleotide^10, 11^. To determine whether two misincorporations on a sequencing read represent two methylation events or a single methylation event which induced multiple misincorporations, we analyzed misincorporation data at endogenously methylated bases. We reasoned that we could determine a minimum separation distance between unique methylation events by plotting the misincorporation rate around endogenously methylated bases. If either SuperScript II or Marathon produced multiple misincorporations for a single methylation site, then there would be an increased misincorporation rate at surrounding un-modified nucleotides. These plots show that SuperScript II produces multiple misincorporations at m^1^G and abasic sites with a complex misincorporation type causing downstream bases to appear highly mutated. Interestingly, SuperScript II encodes m^3^pseudoU as a single point misincorporation. Furthermore, Marathon RT encodes all three modification types as single point misincorporation. This suggests that each unique combination of reverse transcriptase enzyme and modification type may be associated with a unique misincorporation signature. Together, these data suggest that two misincorporations should be separated by at least two non-misincorporated bases, to be confident that these derive from two separate methylated nucleotides, for BASH MaP with SuperScript II.

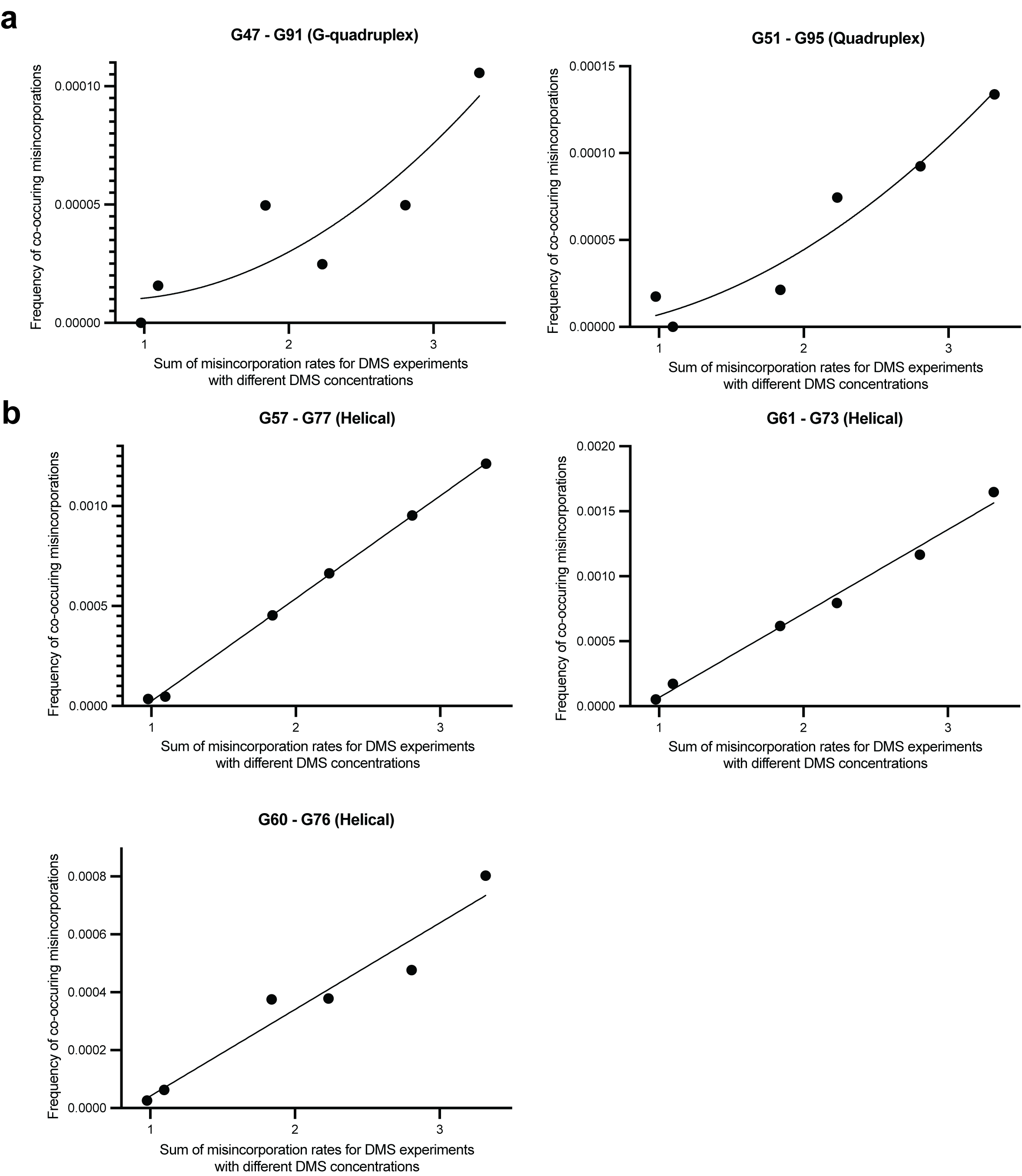

**Supplementary Data Fig. 9 | Discrimination between multi-hit versus single-hit mechanisms for co-occurring misincorporations between G’s in Spinach BASH MaP data**

(**A-B**) BASH MaP was performed on Spinach over a range of DMS concentrations and the frequency of co-occurring misincorporations between specific G residues was plotted. Single-hit versus multi-hit mechanisms for producing co-occurring misincorporations are expected to display different relationships to increasing DMS modification rates. Single-hit mechanisms predict a linear dependence between DMS modification rate and the frequency of co-occurring misincorporations between two locations. In contrast, multi-hit mechanisms predict a quadratic relationship between DMS modification rate and the frequency of co-occurring misincorporations between two locations. Two sets of G-quadruplex G’s display a quadratic relationship while all G’s engaged in Watson-Crick base pairs display a linear relationship.

**Supplementary Table 1 | Spinach G-quadruplex identification false positive rate estimation for various BASH MaP data processing approaches**

| **BASH MaP N7G reactivity (population average) (choose 9 lowest reactivity G’s)** | | | **BASH MaP N7G reactivity (DANCE state 1) (choose 9 lowest reactivity G’s)** | | |
| --- | --- | --- | --- | --- | --- |
| **Sensitivity** | **PPV** | **FPR** | **Sensitivity** | **PPV** | **FPR** |
| 88.88% (8/9) | 88.88% (8/9) | 11.1% (1/9) | 100% (9/9) | 100% (9/9) | 0% (0/9) |

| **BASH MaP N7G reactivity (tertiary folding constraint method)** | | |
| --- | --- | --- |
| **Sensitivity** | **PPV** | **FPR** |
| 77.77% (7/9) | 100% (7/7) | 0% (0/7) |

**PPV = Positive predictive value**

**FPR = False positive rate**
